# Repetition-controllable gain-managed nonlinear fiber amplifier enables ultrashort, multiphoton imaging with reduced photodamage

**DOI:** 10.64898/2026.04.22.720141

**Authors:** James Read, Duanyang Xu, Jikun Yan, Andrew Rawlings, Siddhi Chugh, Mirella Cosma Spalluto, Paul Elkington, Janos M. Kanczler, Simon I.R. Lane, Sumeet Mahajan, Lin Xu

## Abstract

We report a repetition-controllable gain-managed nonlinear fiber amplifier (GMNA) that delivers near-infrared 50-fs pulses with pulse energies up to 150 nJ and a widely tunable repetition rate from 1–20 MHz, while maintaining stable pulse quality across the full range. Using this source, we demonstrate label-free multiphoton imaging—including metabolic autofluorescence (2PF/3PF), second/third-harmonic generation, and Simultaneous Label-free Autofluorescence Multiharmonic (SLAM) microscopy imaging—across live cells, human lung spheroids, and hard tissues. We further assess the impact of laser repetition rate on photodamage at fixed pulse energy, supported by preliminary measurements indicating lower damage at lower repetition rate. Collectively, the compact architecture and repetition-rate agility of the GMNA enable real-time optimization of imaging speed, depth, and sample safety for advanced biological microscopy.

## 2. Introduction

Multiphoton techniques, including fluorescence and harmonic based methods, have gained increasing popularity since their first use in the 1990s [1-3]. The principal multiphoton modalities include two- and three-photon florescence microscopy and second- and third-harmonic generation. These techniques are inherently nonlinear, as excitation occurs through the simultaneous absorption of two or more photons [4]. Such processes require a high photon flux, which is typically achieved at the focal volume of a pulsed laser source. Because signal generation depends nonlinearly on photon flux, out-of-focus signal can be strongly suppressed. As a result, two- and three-photon microscopy provide intrinsic optical sectioning without the need for a confocal pinhole [5]. Furthermore, the absence of out-of-focus excitation reduces photobleaching and photodamage in those regions outside the focal plane. The near-infrared excitation sources commonly used for multiphoton imaging lead to reduced scattering compared to visible wavelengths and typically avoid major absorption peaks of blood and water [6, 7]. Together with the inherent three-dimensional imaging capability, these properties allow for deeper penetration into biological tissues, making multiphoton techniques well suited for imaging thick and highly scattering samples [6].

Two- and three-photon fluorescence have the same contrast generation mechanism as single-photon fluorescence and the same fluorophores can be used for all. The advantages of the multiphoton approach have led two- and three-photon fluorescence microscopy to become the gold-standard technique in several fields, including intravital imaging, deep tissue imaging, and metabolic studies [8-11]. In recent years, three-photon fluorescence has received increasing attention, driven by the growing emphasis on achieving greater imaging penetration [12].

Second- and third-harmonic generation are, on the other hand, complimentary multiphoton label-free techniques that provide contrast based on intrinsic structural properties [1, 3]. Second-harmonic generation (SHG) is a second-order nonlinear process where two low energy photons are combined to produce a single photon at twice the energy. This occurs in non-centrosymmetric structures, for example in biological imaging it is most commonly generated by fibrillar collagen [13], as this is a primary structural component in extracellular matrix across a wide range of biological systems [14-16]. Consequently, SHG is increasingly becoming a widely used imaging modality. Third-harmonic generation (THG) is a third-order nonlinear process where three low energy photons are combined to generate a single photon at three times the energy [17, 18]. In contrast to SHG, the contrast with THG comes from changes in refractive index within interfacial regions, which occurs at cell membranes and other boundary regions within cells and tissues [19].

These techniques can be combined within multimodal imaging systems to simultaneously gather a wide range of contrast mechanisms. In two- and three-photon microscopy, there is a natural spectral separation of signals based on the fact that harmonic emission at exact integer fractions of the excitation wavelength, while fluorescence emission is Stokes shifted to longer wavelengths. As a result, fluorescence and harmonic based signals can be excited and detected concurrently without spectral overlap. Simultaneous excitation and detection of SHG, THG, two- and three-photon auto-fluorescence was demonstrated by You *et al*. in intravital imaging of live mice, and subsequently applied to studies of metabolism, tissue structure and cellular development in mouse skin [8, 20]. This combined multimodal approach was termed Simultaneous Label-free Autofluorescence Multiharmonic (SLAM) microscopy.

SLAM and similar multimodal imaging using multiphoton excitation places specific requirements on the excitation laser source. Near infrared wavelengths above 1µm are typically used to avoid excitation into the deep UV region during three-photon process. However, operation at longer wavelengths presents challenges for intrinsic fluorophores, which generally exhibit low absorption cross sections in this spectral region [21], which can make detecting the generally weak auto-fluorescence difficult. To maximize signal generation, low repetition rate laser sources producing ultrafast pulses are preferred, due to the quadratic and cubic dependence of signal intensity on photon flux for two- and three-photon modalities, respectively. Low repetition rate sources are advantageous because they mitigate average-power-driven effects such as thermal and photochemical damage. However, they can increase the risk of nonlinear photodamage due to high peak intensities, which can be managed through careful control of the average power as highlighted by numerous studies [22-27].

Fiber-based ultrafast laser sources are attractive for biological studies due to their inherent stability and ease of integration, however, conventional fiber oscillators typically operate at fixed repetition rates. Alternative architectures such as Mamyshev oscillators and chirped pulse amplification (CPA) can deliver short pulses with higher energies, but often lack convenient control of repetition rates or have prohibitively complex system designs [28, 29]. Gain-managed nonlinear amplification (GMNA) has recently emerged as a comparatively simple and power-scalable approach for sub-50 fs pulses, achieved by balancing strong nonlinear spectral broadening with longitudinally evolving gain shaping [30-36].

Here, we present a repetition-controllable GMNA 1040 nm excitation source delivering ∼50 fs pulses with pulse energies of 15–150 nJ over a widely tunable repetition-rate range of 1–20 MHz, while preserving comparable spectral/temporal quality across operating points. This repetition-rate agility provides an additional, practically tunable degree of freedom—alongside pulse energy and pulse width—to balance nonlinear signal generation and photodamage/thermal accumulation in multiphoton microscopy. We demonstrate the suitability of the repetition-controllable GMNA laser source for multimodal multiphoton imaging in both complex soft and hard biological tissues. Spheroids made using lung cells and chick bone samples were used as representative biological samples. Organoids and spheroids are an emerging model system in biological research, as they closely recapitulate the structural and functional characteristics of their tissue of origin, including native cellular organization and cell-matrix interactions [37, 38]. However, these self-assembled 3D structures present a significant imaging challenge due to their large size, often exceeding 150 µm, which limits visualization of the central regions using conventional optical imaging methods. Photodamage during multiphoton imaging is also repetition-rate dependent, we assessed it in spheroid samples with the GMNA source. In addition, SLAM imaging was performed on chick bone samples. Bone tissue is particularly challenging to image due to its high degree of mineralization and strong optical scattering. Recent studies have focused on exploiting near-infrared optical windows to enable deeper imaging within such tissues [39, 40]. Thus overall, we demonstrate (i) in-situ dispersion optimization to recover ultrashort pulses through a commercial microscope optical train, (ii) label-free multiphoton imaging across live cells and 3D human lung spheroids, (iii) multimodal SLAM imaging in hard tissue, and (iv) a preliminary repetition-rate–dependent photodamage assessment at fixed pulse energy in 3D spheroids samples.

## 3. Methods - GMNA source and microscope configuration

### Laser architecture

The repetition-controllable GMNA system employs a linear-cavity, semiconductor saturable absorber mirror (SESAM) based, mode-locked Yb-fiber dissipative soliton laser as a seed which delivers 10 ps up-chirped pulses (can be compressed to ∼300 fs) at 1040 nm, with a 3 dB spectral bandwidth of 10 nm, and an average output power of 20 mW at 20 MHz. The seed output is then directed into a fiber-coupled acousto-optic modulator (AOM: AA Optoelectronic MT110-IR20-Fio-PM0.5-J1-A) that selects repetition rates between 1 and 20 MHz. The picked pulse train is subsequently pre-chirped by a diffraction grating-pair based pre-chirp stage to a final ∼ –0.1 ps^2^ of negative group-delay dispersion to compensate the normal dispersion accumulated in the gain fiber and optimize the subsequent nonlinear spectral evolution.

The amplifier stage consists of 3.5 m Yb-doped fiber (PM-YDF 10/125, Coherent) forward-pumped by a 976 nm, 9 W fiber-coupled laser diode through a (2 + 1) × 1 PM pump combiner. The amplifier output is collimated and compressed by a 1000 lines/mm grating-pair compressor with ∼60 % transmission efficiency. The total footprint of the GMNA module is < 0.4 × 0.3× 0.45 *m* .

At the optimum operating point, the system delivers 15–150 nJ pulses tunable from 1 to 20 MHz while maintaining similar spectral and temporal quality across the full range.

### Pulse characterization

Pulse characterization was performed using an optical spectrum analyzer (Yokogawa AQ6370C) and a non-collinear second-harmonic-generation autocorrelator (APE Pulse Check). The output spectra remained smooth and stable across all operating conditions, consistent with GMNA’s gain-shaped nonlinear broadening dynamics [30-36]. Autocorrelation traces confirmed ∼50 fs pulses at 100 nJ for repetition rates from 1 MHz to 20 MHz. At 1 MHz, pulses could also be compressed to ∼50 fs over an energy range of 50– 150 nJ without notable pedestal growth or secondary structures.

### Microscope and detection

The GMNA laser was coupled into a Zeiss LSM 980 microscope equipped with an Examiner Z1 upright stand (Figure 4). The beam was expanded to fill the objective back aperture and scanned using the microscope’s internal optics and galvanometric scanning mirrors. For all experiments, a 20X objective with a numerical aperture (NA) of 1.0 (Zeiss Plan-Apochromat, 421452-9700-000) was used. Live cell imaging was performed using a stage top incubator (OKO-lab H301) providing control of carbon dioxide concentration, temperature and humidity. 3D image stacks were acquired by axial translation of the objective.

**Figure 1.**
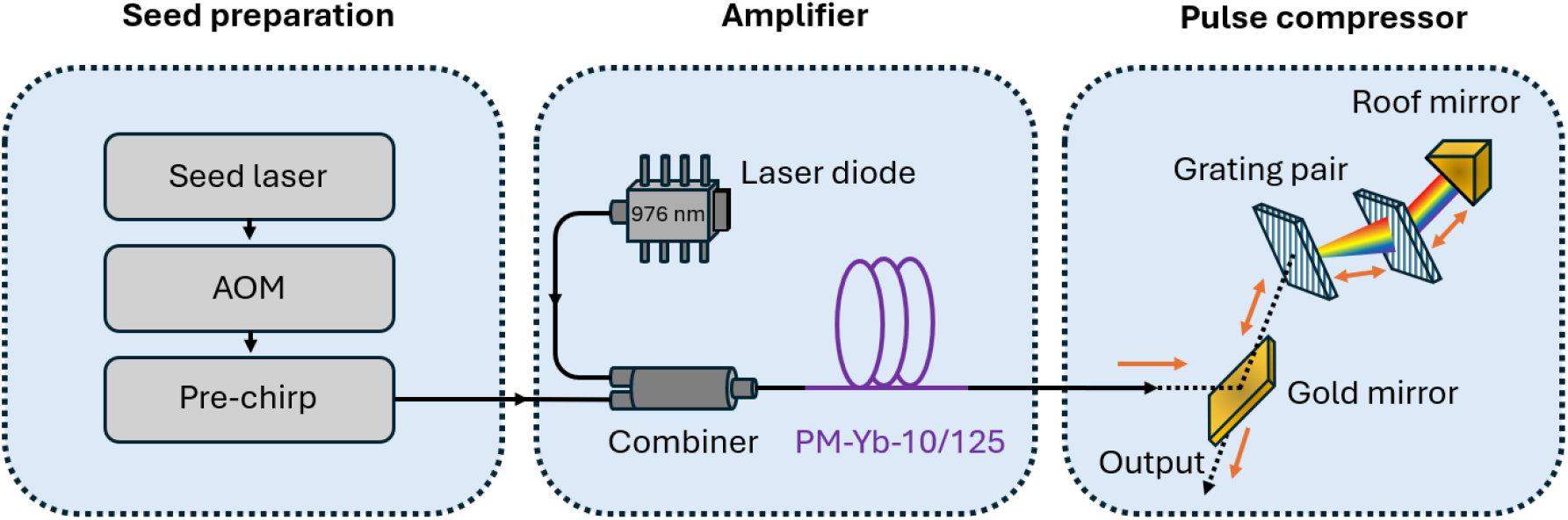
Gain-managed nonlinear fiber amplifier (GMNA) system architecture. A seed laser is pulse-picked by a fiberized acousto-optic modulator (AOM) to select repetition rates, then pre-chirped by a grating pair before amplification in Yb-doped fiber pumped via a pump combiner. The output is collimated and compressed using a grating-pair compressor before delivery to the microscope.

**Figure 2.**
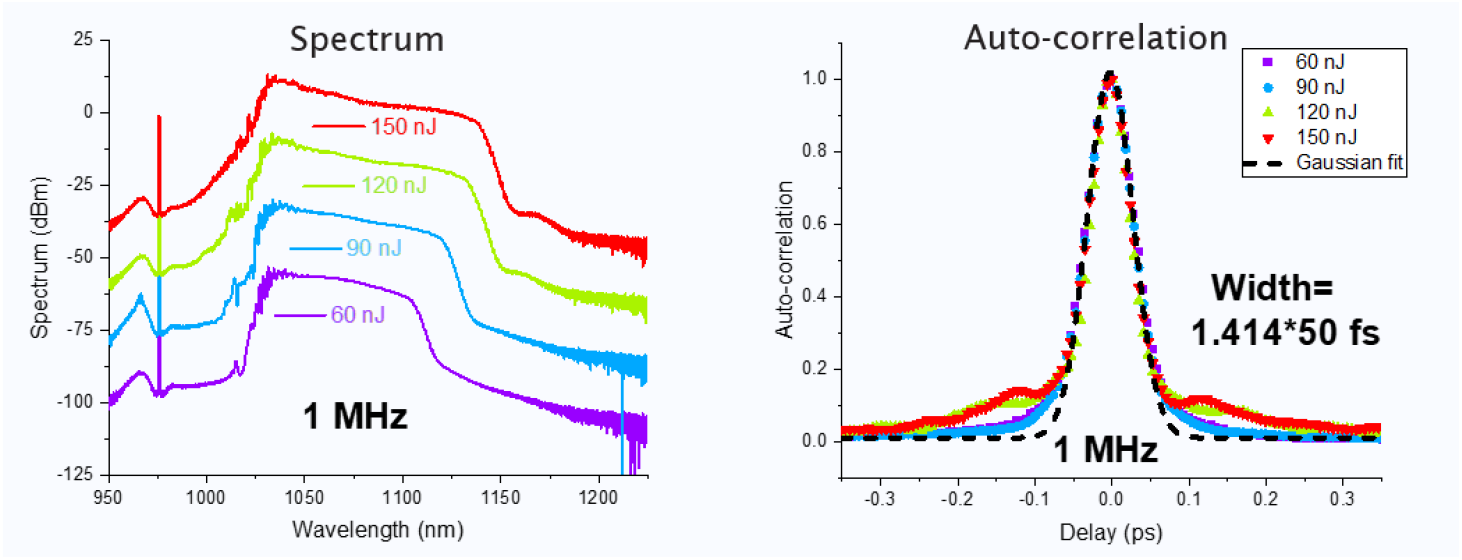
Spectrum & Pulse Characterization at 1 MHz

**Figure 3.**
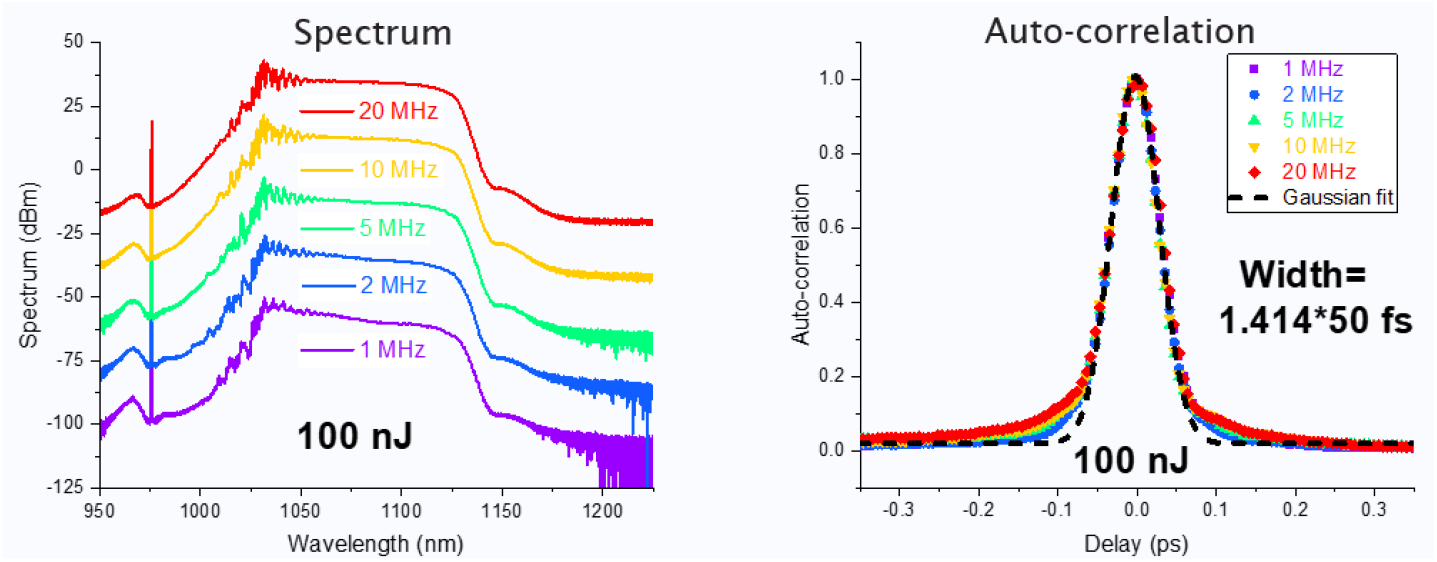
Spectrum & Pulse Characterization at 1-20 MHz at 100 nJ

**Figure 4.**
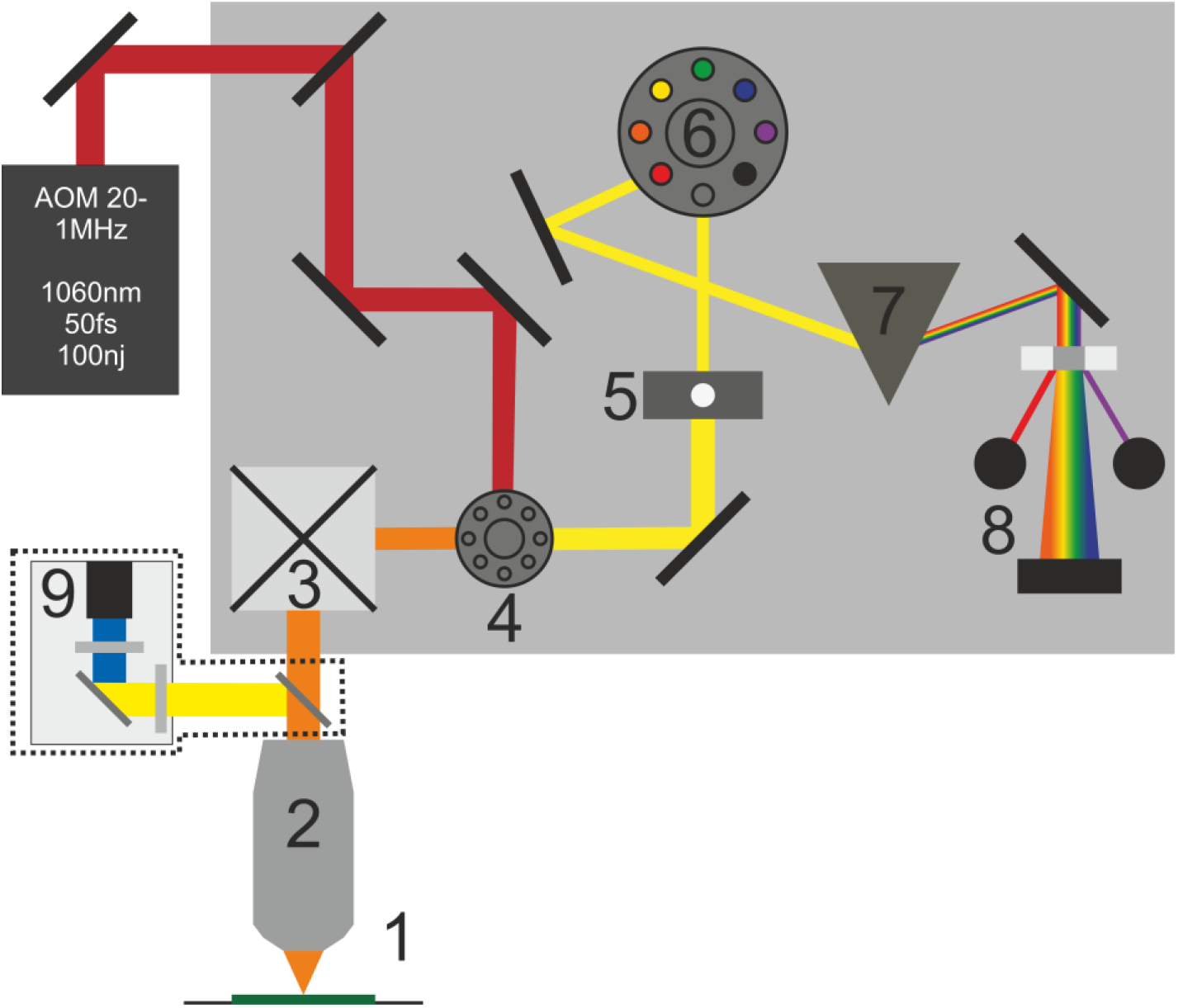
Microscope setup schematic: Zeiss LSM 980 with the GMNA laser, 1 sample; 2, objective; 3, Galvo scanning mirrors; 4, Beam splitter; 5, Pinhole; 6, Secondary beam splitter; 7, Recycling loop (diffraction grating); 8, Confocal detectors; 9, NDD detector with removeable dichroic mirror and bandpass filter (THG only)

For cellular imaging, autofluorescence signals were collected using the internal detectors of the LSM980 microscope, with the detected wavelengths ranges defined by a holographic grating. A multialkali photomultiplier tube (PMT) was used to detect three-photon fluorescence in the 400-490 nm spectral range, while a Gallium arsenide phosphide (GaAsP) PMT was used to collect two-photon fluorescence between 561 nm and 694 nm.

For SLAM imaging, nonlinear signals were collected using both the internal detectors and the epi non-descanned detector. Emission was split into four detection channels. Three internal detectors, including two multi alkali PMT and one GaAsP PMT, were used for signal detection, with spectral separation performed by the diffraction grating. To detect THG signal, a removable 775 nm long-pass dichroic mirror was inserted into the detection path to divert the signal where a 360-10 bandpass filter (Thorlabs, FBH360-10) was used. When the dichroic mirror was removed, the system allowed simultaneous detection of the SHG (500-550 nm), three-photon fluorescence (378-496 nm), and two-photon fluorescence (553-755 nm).

### Spheroids Preparation

BCi-NS1.1 bronchial epithelial cells [41] were employed for spheroid generation. Cells were maintained in PneumaCult™-Ex Plus Medium (STEMCELL Technologies) with routine medium changes and passaging and were not used beyond passage 25.

For spheroid formation, AggreWell™ 400 24-well plates (STEMCELL Technologies) were pre-treated with Anti-Adherence Rinsing Solution according to the manufacturer’s instructions. Cells were seeded at a density of 2.4 × 10^5^ cells per well, corresponding to approximately 200 cells per microwell and incubated for 6 days to promote aggregation and spheroid formation. During this period, partial medium changes were performed using PneumaCult™ Apical-Out (AOAO) Medium, with motile cilia detectable by the end of this aggregation phase.

Following aggregation, spheroids were gently harvested and transferred to Anti-Adherence-treated 24-well flat-bottom plates for continued culture. The medium was replaced every 2 days with PneumaCult™ AOAO Medium. Spheroids were considered differentiated after 1 week in flat-bottom culture and were subsequently used for downstream imaging experiments.

### Imaging preparation – Spheroids

For imaging, the spheroids medium was completely exchanged for phenol red free RPMI (with 10% FBS). The spheroids were then mixed in an equal volume of 0.5% Agarose (made with Hanks’ Balanced Salt Solution) to immobilize the samples. An approximately 30µL droplet was placed at the center of a 35 mm glass bottomed gridded dish (Ibidi, 81168), and additional agarose was placed around the droplet to prevent detachment. RPMI, 2 mL was then added to the dish. This preparation was used consistently for both imaging experiments and photodamage studies.

### Damage and Live dead

Three different repetition rates (1, 5 and 20 MHz) and three different peak powers (104, 52 and 26 kW) were investigated. Photodamage was induced by raster scanning the laser over a 40 × 40 µm^2^ square region centered within the spheroid. This exposure was repeated every 5 µm throughout the spheroid. The pixel dwell time was set to 163.2 µs and all other imaging parameters were kept the same. The methodology of selectively exposing the spheroid core was validated by comparison with damage to a spheroid that was entirely exposed (figure S1).

Following laser exposure, spheroids were returned to AOAO medium and placed in an incubator. After a minimum of 18 hours, samples were removed and the medium was exchanged for RMPI (no FBS) that contained 2 μM calcein AM and 4 μM EthD-1. This was left for 30 minutes before imaging with a widefield fluorescence microscope using appropriate optical filters. Each peak power and repetition rate combination was repeated 4 times.

Imaging analysis was performed using Image J (FIJI). Fluorescence intensities for the Live/Dead stains were quantified by calculating the corrected fluorescence for each channel. This is the measured integrated density of the fluorescence signal from the spheroid corrected for background (this was calculated by measuring the average background intensity and multiplying it by the spheroid ROI area). The ROI was defined using brightfield images to ensure complete coverage of the spheroid. The percentage of cell death was calculated as the ratio of dead fluorescence to total fluorescence and expressed as a percentage. To counter fluorescence settings difference between runs the values from the non-exposed controls from each run was subtracted. A linear mixed-effect model was used; each repeat was treated as a random intercept and repetition rate, peak power, and their interaction as fixed effects. Model fits used maximum likelihood (reml = False). Likelihood-ratio tests compared nested models to access the interaction and main effects. Additionally, each condition was compared to the control normalized (0) using one-sample test and paired t-tests were conducted between each repetition rate within each peak power (paired by repeat). Because multiple comparisons were performed within each peak‐power group, p‐values were adjusted using the Benjamini–Hochberg false discovery rate (FDR) procedure to control the expected proportion of false positives among significant results. For the analysis at fixed average power, we compared normalized percentage death at 26 mW between 5 MHz (104 kW peak) and 20 MHz (26 kW peak) used paired t-test. The statistical analysis was conducted in Python (pandas, stats models, scipy, matplotlib) while Origin (2024b) was used for data plotting.

### Isolation of embryonic chick femur bone sample

A day 18 chick bone sample was taken from fertilized hens’ (Gallus gallus domesticus) eggs from Henry Stewart & Co., Lincolnshire. The eggs were stored at 15±3°c to keep them in stasis. The eggs were slowly brought to room temperature and then placed in a heated humidified environment at 37°c with 60% humidity. The eggs were incubated for 18 days in a horizontal position rotating schedule (1-hour rotations). At day 18 of chick embryo development, the eggshells were cracked open, and embryos were sacrificed by a schedule 1 method and femur bones were isolated. The soft tissue such as adherent muscles and ligaments were carefully removed while preserving the periosteum. These femurs were immediately fixed in 4% PFA [42]. All procedures were conducted with prior ethical approval, in accordance with the Animals (Scientific Procedures) Act 1986, under UK Personal Project License (PPL) number P3E01C456.

## 4. Results

### Live cell autofluorescence with dispersion optimization

Coupling of the GMNA system into the microscope introduced additional material into the optical path, including the relay optics, scan lenses, tube lens, and objective (Figure 4). This could significantly broaden the pulse at the sample plane if not compensated for, leading to reduced multiphoton signal generation.

The pulse duration was measured and optimized using autocorrelation at the output of the GMNA system during its construction, however, direct pulse characterization at the sample plane was not feasible. Consequently, signal generated from the sample was used to compensate for the additional dispersion introduced by the microscope. We performed an in-situ dispersion optimization by adjusting the compressor grating-pair separation distance while monitoring the signal intensity and maximizing the multiphoton signal from a reference sample at fixed average power. 2PF and 3PF signals from live SH-SY5Y cells were captured before dispersion correction (Figure 5 A,B). After optimization, a clear increase in signal intensity and improved structural delineation were observed (Figure 5 C,D), corresponding to an approximate 2.8 times improvement in signal to noise (Figure 5E) ratio. The optimized grating separation corresponds to an effective additional dispersion compensation of approximately 3520 fs^2^, estimated from a 0.55 mm change in grating pair separation. This compensation results in a shorter pulse duration on sample and improved SNR without increasing the average power. These results are consistent with increased peak intensity at constant fluence and highlighted the importance of maintaining ultrashort pulse durations in-situ for efficient multiphoton excitation [8, 43, 44].

**Figure 5:**
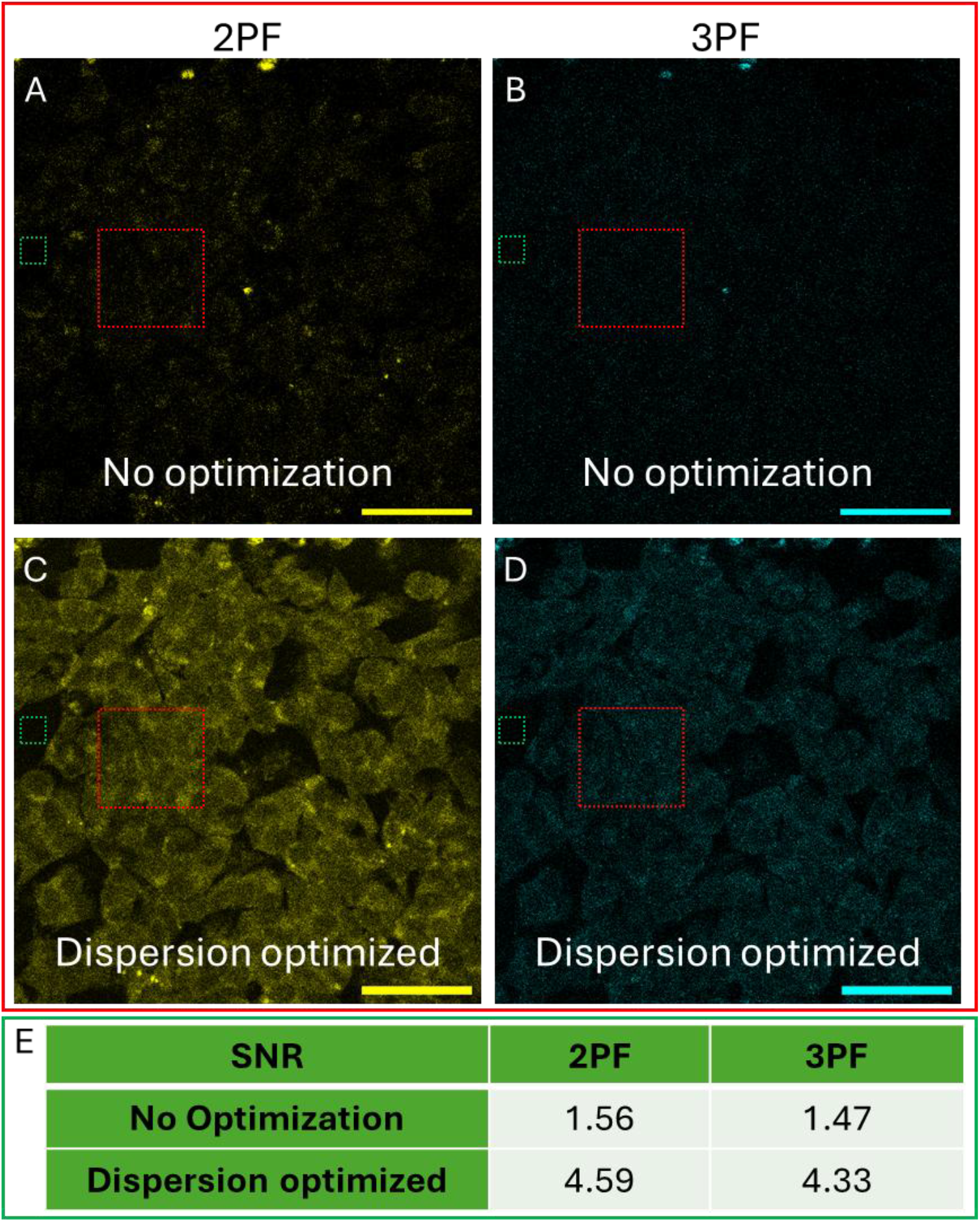
In-situ dispersion optimization to maximize signal. SH-SY5Y 2PF/3PF images with (C and D) and without dispersion optimization (A and B). The red and the green dashed boxes are the ROI’s where the intensity measurements for the background and signal, respectively, were taken from. The table shows the results of the SNR analysis. GMNA: 10 MHz at 100nJ with 50mW average power on sample. 0.96µs pixel dwell time. Scale bar 50µm. 2PF/3PF signals were collected at 553-755 nm and 378-496 nm, respectively.

**Figure 5:**
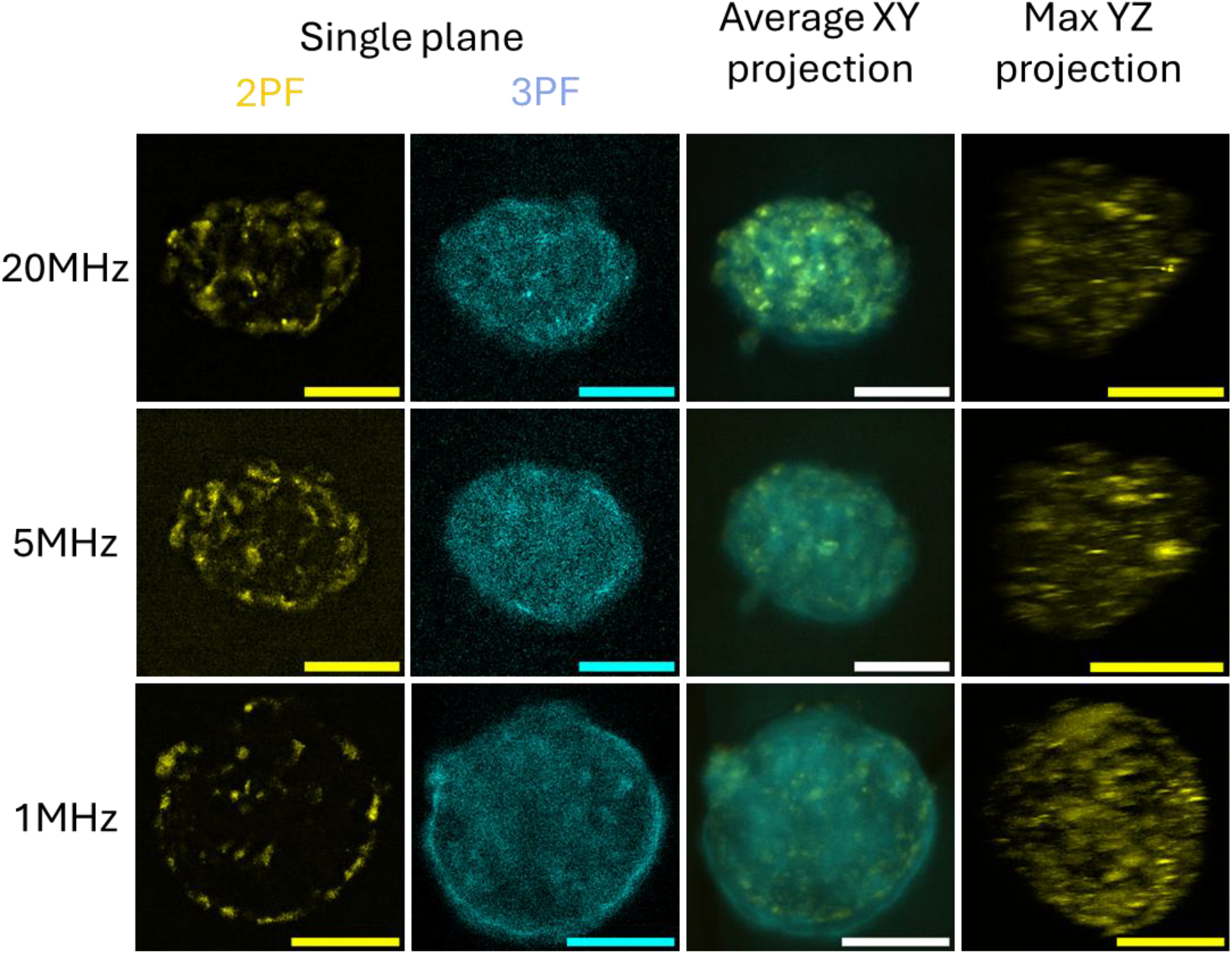
Multiphoton images at different repetition rates. Representative 2PF and 3PF slice frames from different repetition rates with an average projection and a Max YZ projection showing penetration depth. 20 and 5 MHz had a pixel dwell time of 5 µs with 80 mW and 20 mW average power, respectively while 1 MHz had an average power on sample of 6.2 mW with a dwell time of 13.08 µs. 2PF/3PF signals were collected at 553-755 nm and 378-496 nm, respectively. The scale bars for all images are 50 µm.

### Imaging human lung spheroids at variable repetition rates

We next acquired images of live human lung spheroids at repetition rates of 1, 5, and 20 MHz, each operated at ∼100 nJ pulse energy, achieving imaging depths >70 µm with simultaneous excitation of 2PF and 3PF auto-fluorescence using the GMNA laser source. The 2PF and 3PF images collected at 553-755 nm and 378-496 nm, respectively are shown in Figure 6. Small adjustments to dispersion were performed when changing repetition rate; this demonstrates practical, on-the-fly optimization of repetition rate for a given sample while preserving ultrashort pulse quality. At all three repetition rates, both the 3PF and 3PF signals were clearly detectable, with structural features appear throughout the spheroid in both XY and YZ projections, indicating penetration depths of <70µm (Figure 6). The cellular structure of the spheroids is visible under all three repetition rates. This was supported by measuring the signal to noise ratio (SNR) of the two photon autofluorescence which for all repetition rates was >5 even at depths of 70 µm. This high penetration depth was shown by all repetition rates and demonstrates the suitability of the laser for autofluorescence bioimaging of larger samples.

**Figure 6:**
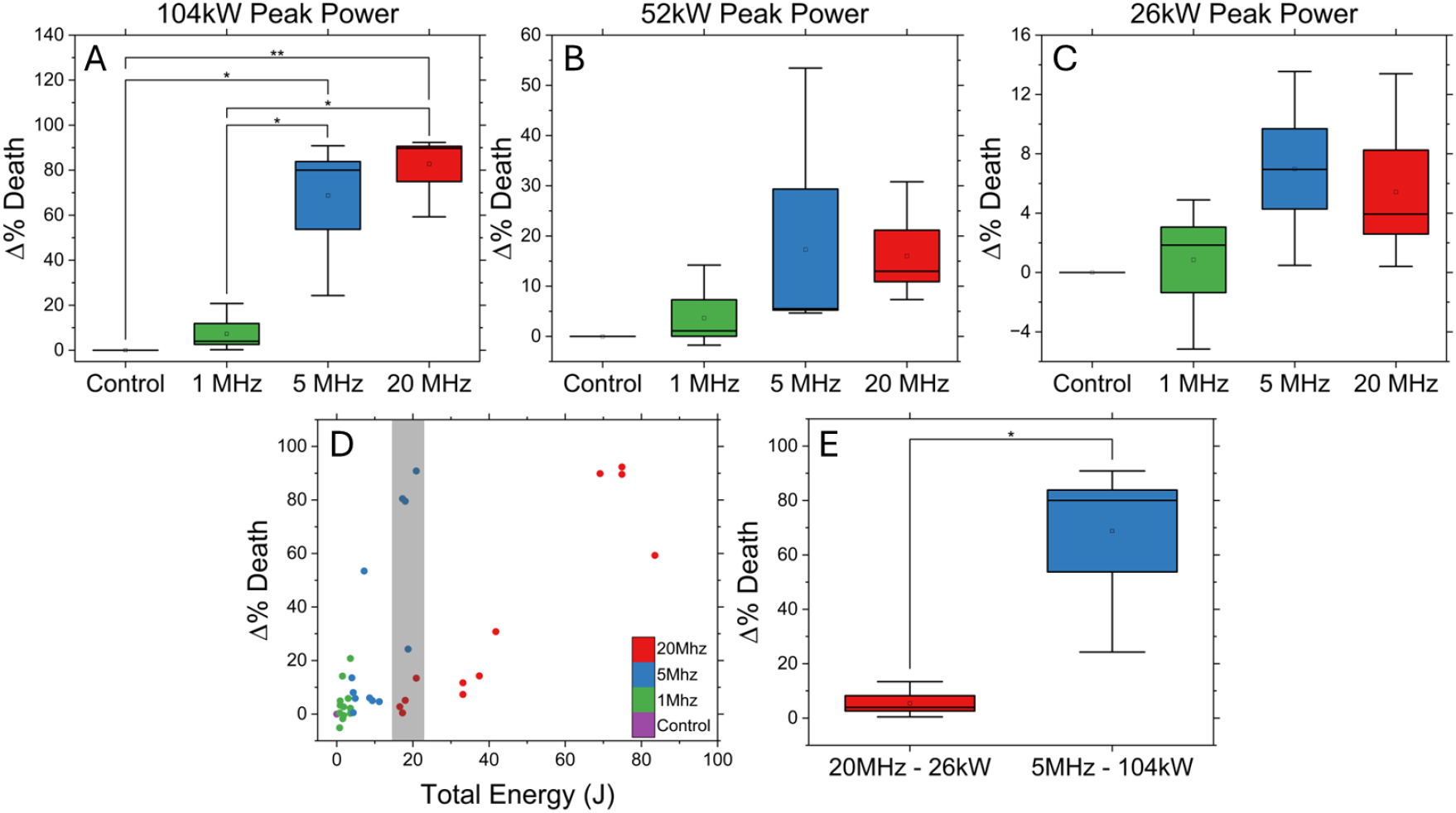
Result of the assessment of the GMNA damage with lung spheroids. Three different peak powers, 104 (A), 52 (B) and 26mW (C) were used with different repetition rates. The positive relationship between the total energy imparted to the spheroid and the damage was plotted (D). The 20 MHz 26 mW peak power and the 5 MHz 104 mW peak power imparted the same total energy this corresponds to the highlighted area in D that was further analyzed in E.

### Repetition-rate–dependent photodamage at fixed pulse energy

To investigate photodamage mechanisms under multiphoton imaging conditions, live dead staining was used to compare percentage cell death at three peak power levels (104, 52 and 26 kW) and three repetition rates (1, 5 and 20 MHz, Figure 7A-C). A linear mixed-effect model was then fitted with each repeat as a random intercept and Repetition Rate, Peak Power, and their interaction as fixed effects. The results show a clear positive correlation between peak power and photodamage, with only 104 kW peak power showing any significant difference to control (figure7A). Lower repetition rates also showed a reduced percentage cell death across all peak powers however significance was only seen at 104 kW between 1-5 MHz and 1-20 MHz and to control at 104 kW for 5 and 20 MHz (Figure 7A). It is highlighted that 1 MHz did not cause significant cell death compared to the control at any peak power.

**Figure 7.**
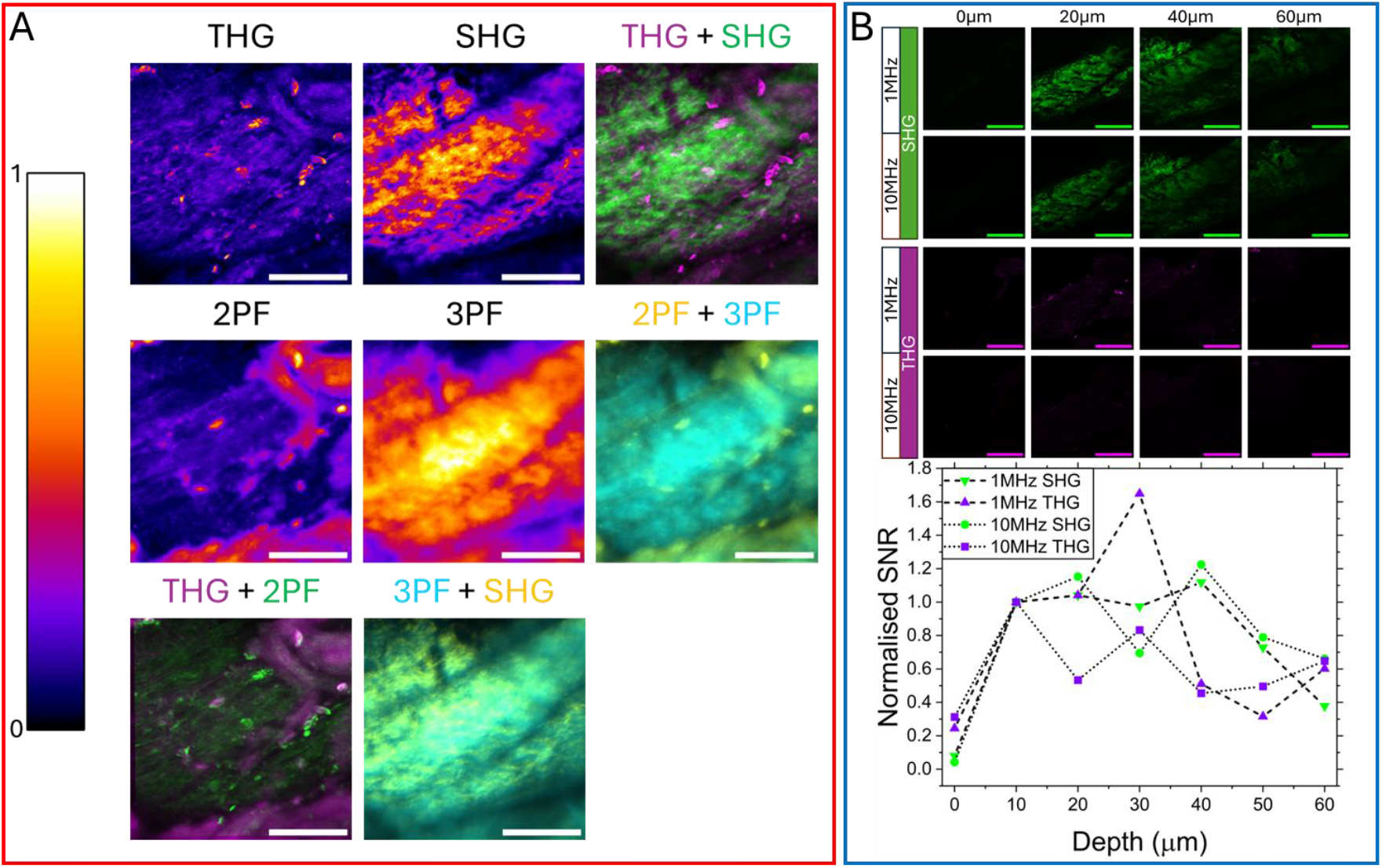
SLAM imaging with GMNA laser. SLAM imaging (THG/SHG/2PF/3PF) was carried out on a day 18 chick bone sample. Part A shows average projections views of all modalities at 1 MHz and selected overlays of multiple modalities showing complementary information. An analysis of imaging depth and signal to noise is shown in part B for 1 and 10 MHz, the peak power was maintained the same, however the pixel dwell time was reduced to ensure the average power was also maintained. Images at 1 and 10 MHz of SHG and THG are shown at 0, 20, 40 and 60 µm depth. Signal to noise was calculated for these images; to allow for simple comparison the values were normalized to the 10 µm depth value. For the 1 MHz images the power at the sample was 6 mW and for the 10 MHz it was 60mW. The pixel dwell time was 5.78 µs for 1 MHz and 0.578 µs for 10 MHz. The scale bar is 50 µm for all images

These observations are consistent with established models where high-repetition-rate excitation increases thermal load and photochemical accumulation between pulses, whereas low-repetition-rate excitation permits thermal diffusion and electronic relaxation [22, 24-27]. The inter-pulse period increases from 50 ns (20 MHz) to 1 μs (1 MHz), providing additional time for heat dissipation and triplet-state relaxation under the low-rep-rate condition. This behavior is commonly associated with reduced photobleaching and photodamage at comparable signal levels, especially in strongly scattering and metabolically active biological specimens [22-27].

Mechanistically, prior studies have reported that (i) damage probability increases with repetition rate at sub-threshold pulse energies due to average-power accumulation; (ii) medium- or low-repetition-rate regimes mitigate cumulative effects; and (iii) distinct damage phenotypes emerge across different repetition-rate regimes. Our preliminary results align with these trends [22, 24-27].

We hypothesized that for a given total energy applied to the sample, a lower repetition rate would be more damaging than a higher one. In our study both the 5 and 20 MHz groups have a test condition where total energy was very close to 20 J (Figure 7D, shaded). Indeed, comparison of percentage cell death at this total energy showed a significant increase in cell death at 5 MHz (Figure 7E, P<0.05). Since the average power between these two groups is the same; it is likely that the total thermal damage is similar, and the significant difference is due to nonlinear damage, i.e. the increased peak power at the low repetition rate is causing the increase in cell death via non-linear damage.

### Hard tissue (bone) SLAM imaging

The full capability of the GMNA laser was explored using Simultaneous Label-free Autofluorescence Multiharmonic (SLAM) imaging. This is a technique that captures unique information from four different techniques using a common illumination wavelength. We tested the suitability of our GMNA laser for SLAM imaging in chick bone at 1 MHz and ∼100 nJ pulse energy (∼6 mW average power on sample). The THG channel, collected at 355-365 nm, highlights refractive-index discontinuities at mineralized interfaces; SHG, collected at 499-552 nm, reports on non-centrosymmetric collagen fibrils; 2PF/3PF, collected at 553-755 nm and 378-496 nm respectively, capture intrinsic fluorophores and higher-order processes. In calcified tissue, SHG texture correlates with collagen orientation and maturation, while THG emphasizes canaliculi and lacunae boundaries and additional interfaces such as cell membranes. From the overlayed images it appears likely that the 3P auto-fluorescence signals come mostly from the collagen based on the strong overlap with SHG while the 2PF is likely from cellular auto-fluorescent compounds such as Flavin Adenine Dinucleotide (FAD) and Nicotinamide Adenine Dinucleotid (NAD). The spatial correspondence between channels further supports their structural specificity[8, 43, 44]. SLAM imaging with a single laser was made possible due to the wavelength range of the excitation generating signals that are compatible with the silicon detector range with all multiphoton techniques used here (2PF, 3PF, SHG and THG).

At 1 MHz chick bone imaging shows penetration to at least 60 µm with clear multimodal contrast. A comparison between 1 and 10 MHz shows that with the same pulse energy the penetration depth is unchanged, to account for changes in average power the pixel dwell for the 10 MHz images was reduced by a factor of ten. A comparison between the two repetition rates is shown in figure 8B, the SHG images reveal that the signal, at 1 and 10MHz is still strong at 60 µm and further depths could be successfully imaged. This is evidenced by the average SNR for SHG, at 1 MHz and 60µm being greater than 20. The THG images have much weaker signals as only parts of the sample had THG active structures. This prevented us from determining the extent penetration depth using THG signal. The SNR comparison between 1 and 10 MHz shows that the rate of decrease over depth is similar for SHG. The zero depth point was complicated by the surface of the sample not being flat. To overcome this the zero depth point was taken to be the frame in which the start of the bulk of signal came into focus at the center of the field of view. Additionally, the signal generating structures were not homogenous at the surface of the bone, this led to weak measured signal (and SNR) at 0µM compared to deeper measurements.

### Discussion

GMNA provides a compact, alignment-light pathway to sub-50-fs, 100-nJ-class pulses with practical repetition-rate agility—capabilities that are typically difficult to obtain simultaneously from oscillators or CPA systems. Recent reports further highlight GMNA’s scalability across wavelengths and regimes (Erbium, sub-MHz, burst mode), suggesting broad applicability for biophotonics and multiphoton imaging [30-36] Our data indicate that repetition-rate tuning is an effective “third knob,” alongside pulse energy and pulse width, to navigate the photodamage–signal trade-off for living samples.

Compared with fixed-repetition-rate sources, our platform allows sample-specific optimization: low repetition rates (e.g., 1–5 MHz) at the same pulse energy can suppress cumulative heating while preserving high per-pulse peak power for deep imaging; higher repetition rates benefit speed where damage tolerance allows. Consequently, lower repetition rates lead to lower photodamage and may be more suitable for live multiphoton imaging applications.

These findings are particularly synergistic with SLAM-style multimodal readouts, which profit from consistent ultrashort pulses to excite multiple endogenous contrast mechanisms simultaneously [8, 43, 44]. It is known that wavelengths over 1000 nm have a lower multiphoton generation efficiency than shorter wavelengths [21]. The resulting loss of signal has been compensated for by high pixel dwell times of 200 µs to 1 ms [45]. Fundamentally it is known that 2P and 3P signals (and photodamage) scales inversely with repetition rate and pulse width [4, 46], indeed it has been shown that for a given SNR a 100 fs pulse showed less photodamage than a 250 fs pulse [47]. The reported GMNA laser system overcomes the challenges of intrinsic weak signal by using short pulses which have a broad bandwidth, with the addition of repetition rate control to balance signal generation and photodamage.

From a laser-engineering perspective, the key advance is the combination of (i) sub-50-fs pulse delivery, (ii) more than 100-nJ-class pulse energy, and (iii) selectable repetition rate over 1–20 MHz in a compact, alignment-light fiber platform. This enables rapid, sample-specific optimization of average power (thermal load) versus per-pulse peak power (nonlinear signal) without changing the imaging wavelength or requiring multiple laser sources. Several past studies looked at laser induced damage on 2D cell culture models[25, 48], this is the first study that looks at photodamage in 3D spheroids. While 2D studies can help inform the 3D studies there are significant challenges unique to 3D systems, the number of times the beam passes though the sample is much higher. In 3D the total sample volume and therefore number of voxels is much greater, meaning the total energy will be higher causing a decrease in pixel dwell time to compensate and therefore having a high peak power to maintain reasonable intensities is vital. This study therefore is an important step towards understanding photodamage in 3D cell models (organoids and spheroids), which increasingly being used across many fields in biomedical research.

## 5. Conclusion

We demonstrate a repetition-controllable GMNA delivering ∼50 fs, up to 150 nJ pulses over 1–20 MHz, enabling label-free multiphoton imaging from soft (spheroids) to hard tissue while maintaining pulse quality across operating points. Preliminary measurements and literature-guided analysis show that, at fixed pulse energy, lowering repetition rate reduces photodamage markers—providing a practical route to deeper, safer imaging. The platform’s simplicity and agility make it a strong candidate for broadly deployable multiphoton imaging microscopes.

## Supporting information

Supplemental doc

## Data/Code availability

Data and analysis scripts will be made available upon reasonable request and deposited in a public repository upon acceptance.

## Acknowledgments

We thank our colleagues at the ORC and in the Biophotonics Group and the Institute for Life Sciences for assistance with sample preparation, imaging, and discussions.

## References

1. Weigelin, B., G.J. Bakker, and P. Friedl, Third harmonic generation microscopy of cells and tissue organization. J Cell Sci, 2016. 129(2): p. 245–55.

2. Denk, W., J.H. Strickler, and W.W. Webb, Two-photon laser scanning fluorescence microscopy. Science, 1990. 248(4951): p. 73–6.

3. Campagnola, P.J. and C.-Y. Dong, Second harmonic generation microscopy: principles and applications to disease diagnosis. Laser & Photonics Reviews, 2011. 5(1): p. 13–26.

4. Zipfel, W.R., R.M. Williams, and W.W. Webb, Nonlinear magic: multiphoton microscopy in the biosciences. Nature Biotechnology, 2003. 21(11): p. 1369–1377.

5. Diaspro, A., et al., Multi-photon excitation microscopy. BioMedical Engineering OnLine, 2006. 5(1): p. 36.

6. Helmchen, F. and W. Denk, Deep tissue two-photon microscopy. Nature Methods, 2005. 2(12): p. 932–940.

7. Key, H., et al., Optical attenuation characteristics of breast tissues at visible and near-infrared wavelengths. Phys Med Biol, 1991. 36(5): p. 579–90.

8. You, S., et al., Intravital imaging by simultaneous label-free autofluorescence-multiharmonic microscopy. Nature Communications, 2018. 9(1): p. 2125.

9. Miller, M.J., et al., Autonomous T cell trafficking examined <i>in vivo</i> with intravital two-photon microscopy. Proceedings of the National Academy of Sciences, 2003. 100(5): p. 2604–2609.

10. Skala, M.C., et al., <i>In vivo</i> multiphoton microscopy of NADH and FAD redox states, fluorescence lifetimes, and cellular morphology in precancerous epithelia. Proceedings of the National Academy of Sciences, 2007. 104(49): p. 19494–19499.

11. Druzhkova, I.N., et al., The metabolic interaction of cancer cells and fibroblasts – coupling between NAD(P)H and FAD, intracellular pH and hydrogen peroxide. Cell Cycle, 2016. 15(9): p. 1257–1266.

12. Horton, N.G., et al., In vivo three-photon microscopy of subcortical structures within an intact mouse brain. Nature Photonics, 2013. 7(3): p. 205–209.

13. Bianchini, P. and A. Diaspro, Three-dimensional (3D) backward and forward second harmonic generation (SHG) microscopy of biological tissues. Journal of Biophotonics, 2008. 1(6): p. 443–450.

14. Bilson, J., et al., Markers of adipose tissue fibrogenesis associate with clinically significant liver fibrosis and are unchanged by synbiotic treatment in patients with NAFLD. Metabolism, 2024. 151: p. 155759.

15. Yeh, A.T., et al., Selective corneal imaging using combined second-harmonic generation and two-photon excited fluorescence. Optics Letters, 2002. 27(23): p. 2082–2084.

16. Williams, R.M., W.R. Zipfel, and W.W. Webb, Interpreting Second-Harmonic Generation Images of Collagen I Fibrils. Biophysical Journal, 2005. 88(2): p. 1377–1386.

17. Squier, J.A., et al., Third harmonic generation microscopy. Optics Express, 1998. 3(9): p. 315–324.

18. Barad, Y., et al., Nonlinear scanning laser microscopy by third harmonic generation. Applied Physics Letters, 1997. 70(8): p. 922–924.

19. Armstrong, J.A., et al., Interactions between Light Waves in a Nonlinear Dielectric. Physical Review, 1962. 127(6): p. 1918–1939.

20. Lee, J.H., et al., Simultaneous label-free autofluorescence and multi-harmonic imaging reveals in vivo structural and metabolic changes in murine skin. Biomed Opt Express, 2019. 10(10): p. 5431–5444.

21. Huang, S., A.A. Heikal, and W.W. Webb, Two-photon fluorescence spectroscopy and microscopy of NAD(P)H and flavoprotein. Biophysical Journal, 2002. 82(5): p. 2811–2825.

22. Masters, B.R., et al., Mitigating thermal mechanical damage potential during two-photon dermal imaging. J Biomed Opt, 2004. 9(6): p. 1265–70.

23. Charan, K., et al., Fiber-based tunable repetition rate source for deep tissue two-photon fluorescence microscopy. Biomedical Optics Express, 2018. 9(5): p. 2304–2311.

24. Macias-Romero, C., et al., Wide-field medium-repetition-rate multiphoton microscopy reduces photodamage of living cells. Biomed Opt Express, 2016. 7(4): p. 1458–67.

25. Hopt, A. and E. Neher, Highly Nonlinear Photodamage in Two-Photon Fluorescence Microscopy. Biophysical Journal, 2001. 80(4): p. 2029–2036.

26. Song, J., et al., SNR enhanced high-speed two-photon microscopy using a pulse picker and time gating detection. Scientific Reports, 2023. 13(1): p. 14244.

27. Gasparoli, F.M., et al., Is laser repetition rate important for two-photon light sheet microscopy? OSA Continuum, 2020. 3(10): p. 2935–2942.

28. Liu, Z., et al., Megawatt peak power from a Mamyshev oscillator. Optica, 2017. 4(6): p. 649–654.

29. Olivier, M., et al., Femtosecond fiber Mamyshev oscillator at 1550 nm. Optics Letters, 2019. 44(4): p. 851–854.

30. Group, W.R. Gain-Managed Nonlinear Amplifier (GMNA) guide. [cited 2025.

31. Stoliarov, D., et al., Gain-managed nonlinear amplification of ultra-long mode-locked fiber laser. Optics Express, 2023. 31(26): p. 43427–43437.

32. Tomaszewska-Rolla, D., et al., A comparative study of an Yb-doped fiber gain-managed nonlinear amplifier seeded by femtosecond fiber lasers. Scientific Reports, 2022. 12(1): p. 404.

33. Chen, K., et al., Compact all-PM fiber gain-managed nonlinear amplification system based on a stretched-pulse oscillator. Optics Letters, 2025. 50(7): p. 2129–2132.

34. Xia, T., et al., All-fiber integrated gain-managed nonlinear amplification system delivers ultrafast lasers with 3.6 MW peak power and 45 fs pulse duration. Optics & Laser Technology, 2025. 181: p. 111627.

35. Maghsoudi, A., et al., 50-fs pulse bursts via gain-managed nonlinear amplification. EPJ Web Conf., 2024. 307: p. 02049.

36. Sobon, G. and M. Krakowski, Gain-managed nonlinear amplification in an Erbium-doped fiber. 2024, Optica Open.

37. Lancaster, M.A. and J.A. Knoblich, Organogenesis in a dish: Modeling development and disease using organoid technologies. Science, 2014. 345(6194): p. 1247125.

38. Antoni, D., et al., Three-Dimensional Cell Culture: A Breakthrough in Vivo. International Journal of Molecular Sciences, 2015. 16(3): p. 5517–5527.

39. Mi, C., et al., Bone disease imaging through the near-infrared-II window. Nat Commun, 2023. 14(1): p. 6287.

40. Mateos, S., et al., Instantaneous In Vivo Imaging of Acute Myocardial Infarct by NIR-II Luminescent Nanodots. Small, 2020. 16(29): p. 1907171.

41. Walters, M.S., et al., Generation of a human airway epithelium derived basal cell line with multipotent differentiation capacity. Respir Res, 2013. 14(1): p. 135.

42. Smith, E.L., et al., The Effects of 1α, 25-dihydroxyvitamin D3 and Transforming Growth Factor-β3 on Bone Development in an Ex Vivo Organotypic Culture System of Embryonic Chick Femora. PLOS ONE, 2015. 10(4): p. e0121653.

43. Wang, G., S.A. Boppart, and H. Tu, Compact simultaneous label-free autofluorescence multi-harmonic microscopy for user-friendly photodamage-monitored imaging. J Biomed Opt, 2024. 29(3): p. 036501.

44. De la Cadena, A., et al., Simultaneous label-free autofluorescence multi-harmonic microscopy driven by the supercontinuum generated from a bulk nonlinear crystal. Biomedical Optics Express, 2024. 15(2): p. 491–505.

45. Tu, H., et al., Stain-free histopathology by programmable supercontinuum pulses. Nature Photonics, 2016. 10(8): p. 534–540.

46. Xu, C., et al., Multiphoton fluorescence excitation: new spectral windows for biological nonlinear microscopy. Proceedings of the National Academy of Sciences, 1996. 93(20): p. 10763–10768.

47. Débarre, D., et al., Mitigating Phototoxicity during Multiphoton Microscopy of Live Drosophila Embryos in the 1.0–1.2 µm Wavelength Range. PLOS ONE, 2014. 9(8): p. e104250.

48. Baumgart, J., et al., Repetition rate dependency of reactive oxygen species formation during femtosecond laser-based cell surgery. J Biomed Opt, 2009. 14(5): p. 054040.

