## Supplemental doc for "Repetition-controllable gain-managed nonlinear fiber amplifier enables ultrashort, multiphoton imaging with reduced photodamage"

| | 20MHz | 5MHz | 1MHz |
|---|---|---|---|
| 2PF | 11.42 | 4.59 | 7.57 |
| 3PF | 2.06 | 2.03 | 2.09 |

**Supplementary table 1: The signal to noise ratio (SNR) for both two and three photon auto-fluorescence for each repetition rate was calculated from the single frame images of the live spheroids (figure 6). Three ROIs were drawn onto each spheroid; the average intensity was calculated. The background standard deviation was measured, and the SNR was calculated by $SNR = \frac{Signal_{mean}}{Background_{Std\ Dev}}$. The SNR of the 2PF did decrease with lowing repetition rate, this correlation is due to the decreasing average power. The 5MHz is lower than 1MHz this due to the 5MHz being the same spheroid as the 20MHz example, as the 20MHz measurement was taken first, bleaching has taken place.**

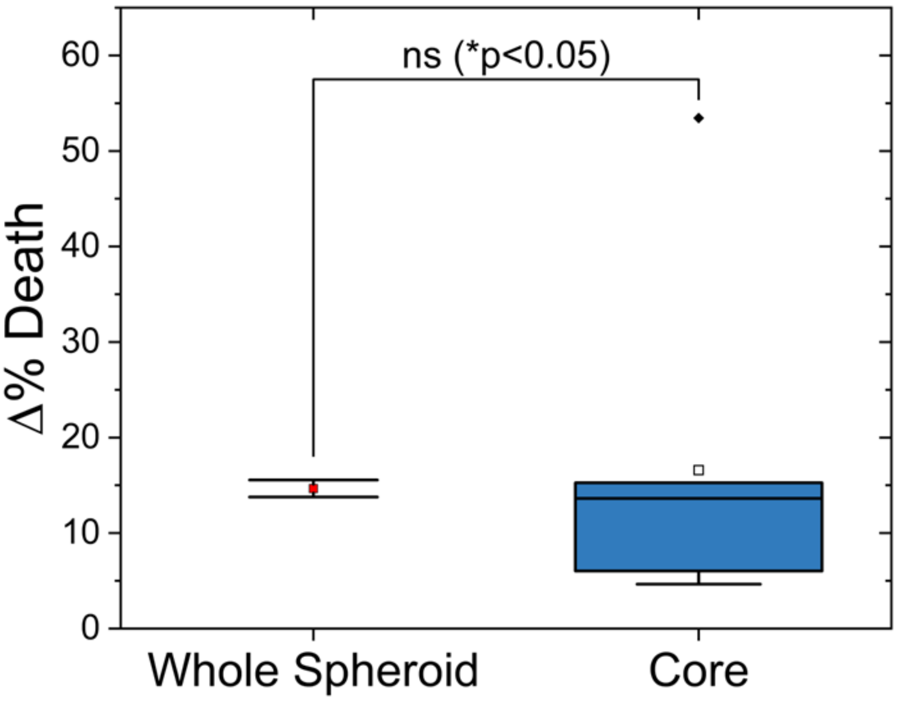


Supplementary figure 1: As explained in the method section, for the spheroid 3D damage study only the core was exposed. This is different for imaging experiments so to validate of this method; an entire spheroid was exposure using the same parameters as one of the experiments. A one sample t-test was used to see if there was any significant difference. The results show there is no significant difference p= 0.22 which is greater than 0.05.
